# Selection of Sites for Field Trials of Genetically Engineered Mosquitoes with Gene Drive

**DOI:** 10.1101/2021.04.28.441877

**Authors:** G.C. Lanzaro, M. Campos, M. Crepeau, A. Cornel, A. Estrada, H. Gripkey, Z. Haddad, A. Kormos, S. Palomares, W. Sharpee

## Abstract

Novel malaria control strategies using genetically engineered mosquitoes (GEMs) are on the horizon. Population modification is one approach wherein mosquitoes are engineered with genes rendering them refractory to the malaria parasite coupled with a low-threshold, Cas9-based gene drive. When released into a wild vector population, GEMs preferentially transmit these beneficial genes to their offspring, ultimately modifying a vector population into a non-vector one. Deploying this technology awaits evaluation including ecologically contained field trials. Here, we consider a process for site selection, the first critical step in designing a trial. Our goal is to identify a site that maximizes prospects for success, minimizes risk, and serves as a fair, valid, and convincing test of efficacy and impacts of a GEM product intended for large-scale deployment in Africa. We base site selection on geographical, geological, and biological, rather than social or legal, criteria. We recognize the latter as critically important but not preeminent. We propose physical islands as being the best candidates for a GEM field trial and present an evaluation of 22 African islands. We consider geographic and genetic isolation, biological complexity, island size, topography, and identify two island groups that satisfy key criteria for ideal GEM field trial sites.

## 1. Introduction

We present a framework employed by the University of California Irvine Malaria Initiative (UCIMI) for the selection of sites to conduct field trials of a genetically engineered mosquito (GEM) with gene drive. These GEMs are designed to offer safe, cost-effective, and sustainable malaria control in sub-Saharan Africa. This will be achieved using a population modification strategy [1] wherein parasite blocking effector genes are engineered into vector mosquitoes, rendering them incapable of transmitting the parasite [2]. An essential GEM component is an efficient gene drive [3] which serves two critical purposes: to establish the effector genes at high frequency in the mosquito population at the immediate release site and to facilitate its spread into neighboring populations via normal mosquito dispersal and gene flow. This GEM is designed to eliminate the malaria parasite without eliminating the mosquito.

Achieving malaria control on a large spatial scale, requires a so-called low-threshold gene drive; meaning one with a maximum capability for spreading across the environment (invasiveness). Henceforth, when we refer to a GEM, we mean a mosquito engineered with anti-*Plasmodium* effector genes and a low threshold, highly invasive gene drive. This is the GEM that UCIMI aims to evaluate in a field trial.

Our primary goal for a site selection process is identification of a site that maximizes the prospects for success, minimizes risk, and serves as a fair, valid, and convincing test of the efficacy and impacts of a GEM product intended for large-scale deployment in sub-Saharan Africa. The purpose of the field trial itself is to describe the behavior of a GEM when introduced into a natural population of a target species, in this case *Anopheles gambiae* and/or its sister species *Anopheles coluzzii*.

A multi-phase pathway for the development and evaluation of GEMs has been proposed by the World Health Organization (WHO) [4]. This protocol has been widely endorsed [5, 6] and serves as the foundation for the framework described here. PHASE 1 of the WHO pathway includes design and construction of the GEM product and initial evaluation of its efficacy. This evaluation assesses the phenotype generated by the transgenes, transgene inheritance (especially as it relates to the efficiency of the gene drive component), the stability of the construct over time, and a rudimentary evaluation of overall fitness [3, 7]. GEM products that show promise then move into PHASE 2 field trials with a strong emphasis on containment.

Early guidelines recommended that initial tests be conducted in large, artificially contained greenhouse-like cages designed to simulate natural conditions [8–11]. Data generated in such caged environments are limited in several important ways: they do not allow analysis of community and ecosystem-level interactions in any meaningful sense, they cannot replicate food web structure, and they do not permit examination of ecological phenomenon (e.g., dispersal) across spatial scales [12, 13]. Critically, experiments conducted in artificial environments often yield highly replicable, but spurious results [14]. These limitations were recognized in later guidelines and the use of artificially contained environments is now suggested as optional, unless required by regulatory authorities [15, 16].

A different strategy that has been proposed for dealing with containment is to conduct field trials in a stepwise fashion with early trials using high-threshold drives, such as split-drive systems which have limited invasiveness and are therefore self-contained [17, 18]. Threshold-dependent drives have their place in controlling vectors on a small spatial scale, such as in urban settings [19]; however, deploying a high threshold drive to achieve malaria control at the scale of continental Africa is not feasible [15].

From our perspective, conducting trials in large cages or with high-threshold drives does not satisfy our goal that field tests be valid and convincing. Therefore, we propose to use ecologically confined PHASE 2 field trials in their place. The issue of containment can be mitigated by selecting the appropriate site.

The first consideration in the selection of a GEM field site should be based on defining biological and physical characteristics that would make a site ideal, or as near to ideal, as possible [8, 11, 20]. Ethical, social, and legal issues are critically important, and no field test can be undertaken before these are addressed [21–23]. However, valuable resources, relationships, and infrastructure are best developed at a site that has been determined to be scientifically suitable. Here we describe a set of criteria that may be applied to a thoughtful consideration and assessment of potential field trial sites. When completed, this framework should provide a cogent justification for why a particular site was selected for GEM testing.

Ecologically confined field sites offer geographical, environmental, and/or biological confinement [4]. Physical islands have been suggested as ideal for conducting GEM field trials [6, 11]. In addition to containment, islands have numerous characteristics that favor their use as GEM field trial sites, including relatively small size, distinct boundaries, simplified biotas, and relative geological youth. These features led to the development of the “Dynamic Equilibrium Theory of Island Biogeography” [24–27], which we rely on to inform our assessment of the advantages of island over mainland sites for the evaluation of GEM.

## 2. Materials and methods

### (a) Selection of candidate island sites

Site selection was initiated with the identification of all potential island sites, which we define broadly as any island associated with the continent of Africa (Figure 1).

**Figure 1.**
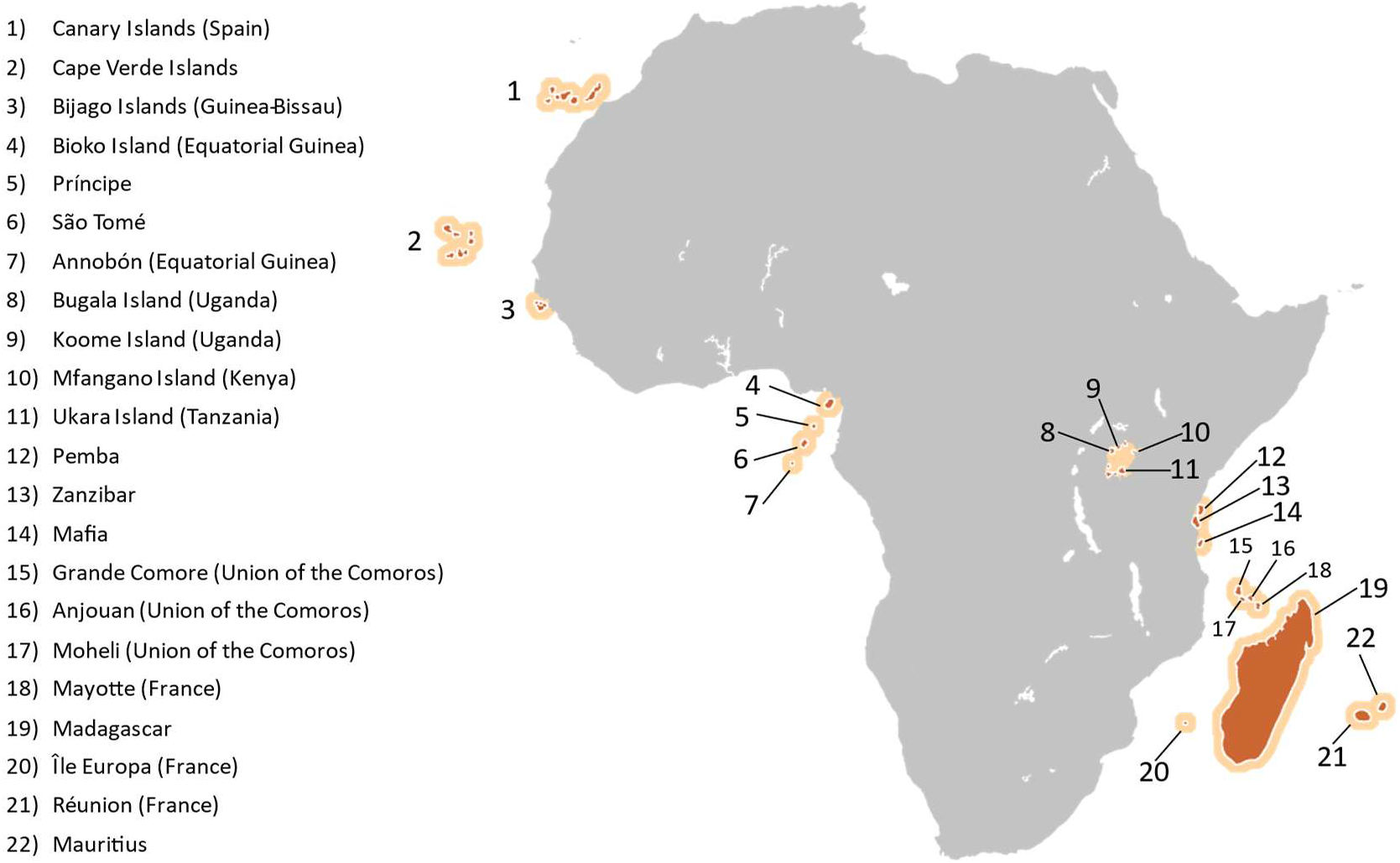
African islands and island groups considered potential field sites for genetically engineered mosquitoes for malaria eradication.

Data for each site was obtained from published sources except for some genetic data which was generated *de novo* by us. These data were used to inform the suitability of potential sites by determining if they meet the set of criteria listed in Box 1. This information includes descriptions of entomological, genetic, geographic, and geophysical features of the sites and mosquito populations therein.

#### BOX 1. Criteria for the selection of field sites.

| **BOX 1.** Criteria for the selection of field sites. |  |
| --- | --- |
| **Primary Criteria** | **Rationale** |
| 1. Presence of *Anopheles gambiae*/*A. coluzzii* | Major vector; target of GEM production |
| 2. Geographic isolation | Containment of GEMs |
| 3. Genetic isolation | Containment of transgene constructs |
| 4. Genetic diversity | Potential detriment to GEM function |
| 5. Island size | Feasibility vs. validity |
| 6. Topography | Evaluation of GEM dispersal capacity |
| 7. Anopheline species richness | Logistics; HGT*; confound epi. endpoints |
| **Other considerations** | **Rationale** |
| 1. Insecticide susceptibility/resistance | Match GEM to indigenous mosquitoes |
| 2. *Plasmodium* prevalence | Estimation of epidemiological impact |
| 3. Presence of endangered species | Potential for negative GEM interactions |
| 4. Travel feasibility | Operational logistics and cost |

\*HGT=horizontal gene transfer

### (b) Measuring Island Geographic Isolation

Geographic isolation for each island was defined using three methods, all reported in Table 1. The first is simply the geographic distance to the nearest mainland. Distances for individual islands were calculated as the shortest great circular distance between an island’s mass centroid and the mainland coast. For archipelagos, distances from the nearest island to the mainland were used [28]. Distance to the mainland for each Lake Victoria island and for Annobón was determined using Google Earth’s distance and area measuring tool. The two closest points on the mainland and island shores were used as measuring points. The significance of distance to mainland is that the nearest mainland is assumed to be the richest gene pool and the source of populations on the islands [28, 29].

**Table 1.** Bioclimatic and isolation index values used for the evaluation of potential island field sites. DD = decimal degrees; UNEP = United Nations Environment Programme; SLMP = Surrounding Landmass Proportion; GMMC = Glacial Maximum Mainland Connection a proxy variable for island geological history which indicates whether an island was connected to the mainland during the Last Glacial Maximum (LGM) (1 = true and 0 = false); - refers to missing or incomplete data. Additional data sources: Bugala:[30–33]; Mfango [34]; Ukara: [35–37]; Koome [38–42]

| Island | Archipelago | Island type | Area (km<sup>2</sup>) | Distance to mainland (km) | UNEP Isolation Index | SLMP | GMMC | Elevation max (m) |
| --- | --- | --- | --- | --- | --- | --- | --- | --- |
| Canary Islands | Canary Islands | Oceanic | 7509.66 | 116.63 | 30.4 | 0.812 | 0 | 3705 |
| Cape Verde | Cape Verde Islands | Oceanic | 4088.52 | 586.53 | 55 | 0.466 | 0 | 2813 |
| Bijagos | Bijagos Islands | Continental | 1944.72 | 0.83 | 10.8 | 1.123 | 1 | 59 |
| Bioko | Cameroon Line | Continental | 1950.46 | 73.03 | 17 | 1.148 | 1 | 3011 |
| Annobón | Cameroon Line | Oceanic | 15.7 | 350 | 45 | - | 0 | 587 |
| São Tomé | Cameroon Line | Oceanic | 854.8 | 283.63 | 39 | 0.753 | 0 | 1977 |
| Príncipe | Cameroon Line | Oceanic | 143.16 | 221.72 | 39 | 0.86 | 0 | 934 |
| Bugala | Lake Victoria | Lacustrine | 296 | 3.7 | 5.487 | - | 1 | 160 |
| Koome | Lake Victoria | Lacustrine | 100 | 14.3 | 10.688 | - | 1 | 180 |
| Mfangano | Lake Victoria | Lacustrine | 66 | 7.4 | 10.414 | - | 1 | 551 |
| Ukara | Lake Victoria | Lacustrine | 80 | 22.8 | 15.623 | - | 1 | 162 |
| Pemba | Zanzibar | Continental | 987.08 | 68.6 | 31.231 | 1.178 | 0 | 149 |
| Zanzibar | Zanzibar | Continental | 1591.5 | 50.9 | 17 | 1.337 | 1 | 133 |
| Mafia | Mafia | Continental | 443.24 | 36.06 | 29.432 | 1.194 | 1 | 66 |
| Grand Comore | Comoros | Oceanic | 1021.61 | 307.45 | 49 | 0.736 | 0 | 2368 |
| Moheli | Comoros | Oceanic | 212.09 | 340.61 | 49 | 0.754 | 0 | 793 |
| Anjouan | Comoros | Oceanic | 432.08 | 417.78 | 49 | 0.706 | 0 | 1591 |
| Mayotte | Comoros | Oceanic | 371.42 | 490.16 | 47 | 0.669 | 0 | 636 |
| Madagascar | Madagascar | Oceanic | 590547.4 | 780.51 | 58 | 0.46 | 0 | 2876 |
| Ile Europa | French Territory | Oceanic | 32.64 | 492.95 | 67.941 | 0.736 | 0 | 20 |
| Réunion | Mascarene Islands | Oceanic | 2512.65 | 1699.32 | 73 | 0.467 | 0 | 3066 |
| Mauritius | Mascarene Islands | Oceanic | 1868.44 | 1874.49 | 87 | 0.399 | 0 | 816 |

A second metric is the United Nations Environment Programme (UNEP) Isolation Index, which is calculated as “the sum of the square roots of the distances to the nearest equivalent or larger island, the nearest group or archipelago, and the nearest continent [43].” The higher the value, the more geographically isolated the island is.

The third isolation index is Surrounding Land Mass Proportion (SLMP) where the isolation of the focal island is proportional to the area of the surrounding landmass [28]. SLMP is calculated as the sum of the proportions of landmass within buffer distances of 100, 1000, and 10,000 km around the island perimeter. SLMP accounts for the coastline shape of large landmasses by considering only regions that extend into the measured buffers. SLMP values for the Canary Islands, Cape Verde Islands, and Bijagós Islands were represented as the average of all islands in their respective archipelagos [28]. SLMP is a preferred index for analysis of species variation on a focal island. The equilibrium theory of island biogeography supports this index as individual islands may act as stepping-stones for species dispersal and establishment, which this index accounts for by shortening the distance between an island and potential source populations [26]. A larger SLMP value indicates that an island is surrounded by more landmass. For this study, we are focusing on islands with a lower SLMP value since these islands will have less surrounding landmass which could facilitate mosquito dispersal into or out of the target island.

### (c) Island size and topography

Island size information, presented in Table 1 as area, is taken from the publication by Weigelt et al. [28]. They describe island size by using the Database of Global Administrative Areas (GADM) to obtain high-resolution island polygons. Area was calculated for each GADM polygon in a cylindrical equal area projection. Areas for archipelagos (Canary Islands, Bijagos, Cape Verde) were reported here as the sum of all islands in each archipelago [28]. The area for Annobón was obtained from the United Nations Environmental Programme [43]. The areas for the Lake Victoria islands (excluding Koome) were taken from the literature [33–35] and the area for Koome Island was approximated using Google Earth’s distance and area measuring tool.

Elevation maximum and minimum of each island were obtained from the AW3D30 Global Digital Surface Model of the Japan Aerospace Exploration Agency [44]. GeoTIFF files were downloaded, and the highest elevation of each island/archipelago was identified. Island topography was further described using the United States National Aeronautics and Space Administration (NASA) 90-m resolution elevation data from the Shuttle Radar Topography Mission (SRTM) 90m Digital Elevation Model database [45]. In this case, altitude and magnitude of steepest gradient measurements were used to generate heat maps as graphic descriptors of topography.

### (d) Population genomics analyses

We conducted a comparative genomics analysis of mainland and island populations of the two target species. The locations and sample sizes per site are provided in Figure 2. In total, 420 individual *Anopheles gambiae* and *A. coluzzii* genome sequences were analyzed in this study. The UC Davis Vector Genetics Laboratory (VGL) generated 167 genomes (Supplemental Table 1). In addition, 196 genomes were obtained from the *Anopheles gambiae* 1,000 Genome Project phase 2 [46] and 57 were taken from a published Lake Victoria islands study [47].

**Figure 2.**
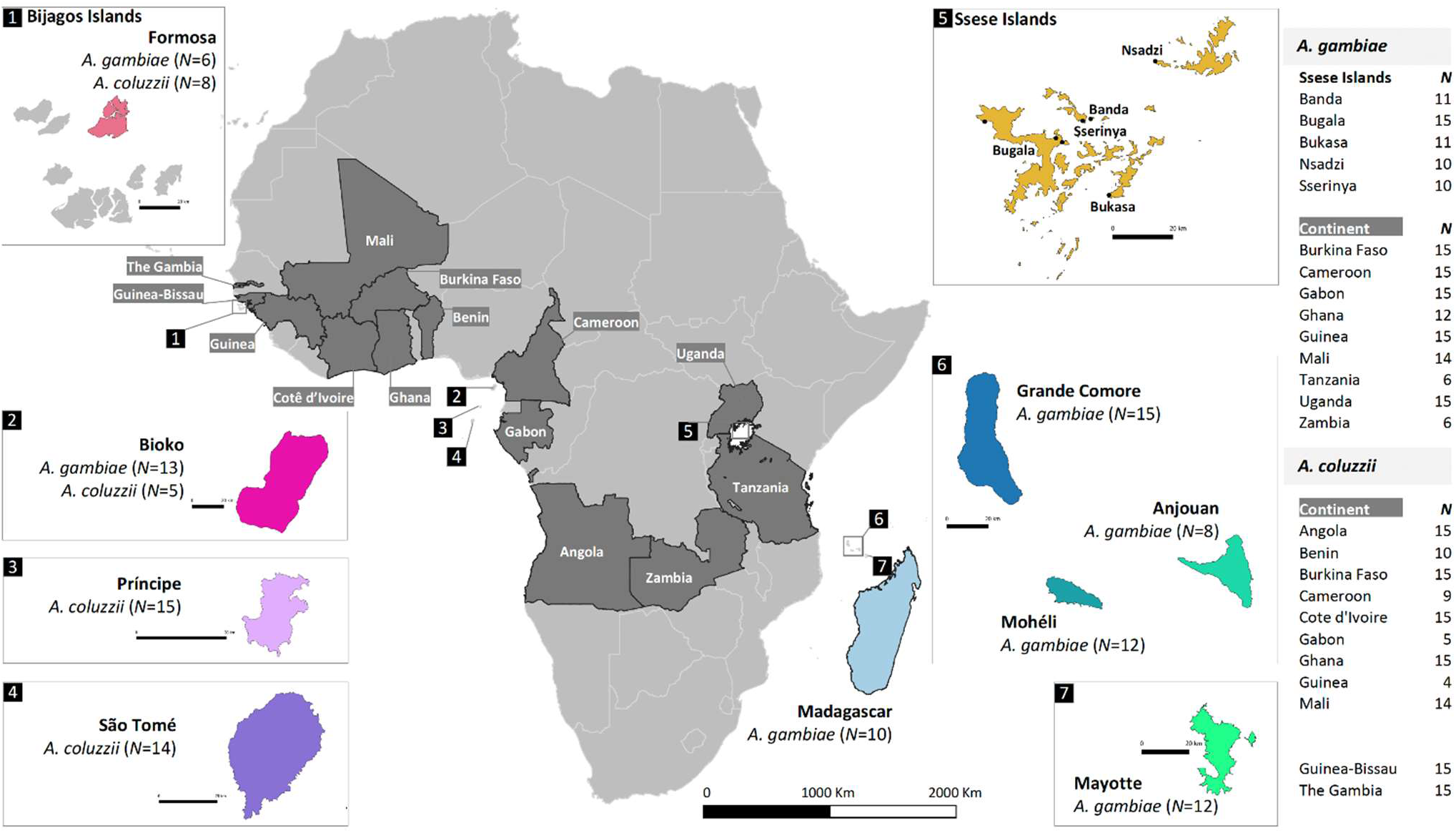
Study sampling locations. Samples of *A. gambiae* and *A. coluzzii* were from 12 countries (dark shade) in continental Africa: Angola, Benin, Burkina Faso, Cameroon, Cotê d’Ivoire, Gabon, Ghana, Guinea, Tanzania, Uganda and Zambia. Populations from Guinea-Bissau and The Gambia (dark shade) were included with no species assignment. Table on the right displays the number of samples for each of the mainland populations. The insert maps show African islands sampled in this study: 1) Formosa islands within Bijagos archipelago; 2) Bioko, 3) Príncipe and São Tomé, 4) islands in the Gulf of Guinea, 5) five islands in Ssese islands in Lake Victoria in Uganda, 6) in the Comoros (Anjouan, Mohéli and Grande Comore), and 7) Mayotte. Madagascar island is shown in the main map. The number of samples included for each island is shown in parenthesis. Insert maps contain a scale of 20km length.

Individual mosquito DNAs from the VGL samples were extracted using a Qiagen Biosprint (Qiagen, Hilden, Germany) following our established protocol [48]. 10 ng of genomic DNA was used for individual libraries using a KAPA HyperPlus Kit (Roche Sequencing Solutions, Indianapolis, Indiana, USA), as described in Yamasaki et.al [49]. Sequencing was performed on an Illumina HiSeq 4000 instrument (Illumina, San Diego, California, USA) at the UC Davis DNA Technologies Core facility. Methods used for genome sequencing of individuals from other sources are described elsewhere [47, 50].

Demultiplexed raw reads of VGL samples were filtered and trimmed using Trimmomatic v0.36 [51] and saved as FastQ files. Sequences from Ag1000G and Lake Victoria study [47] were downloaded and converted to FastQ files using BEDTools v.2.2 [52]. All specimens were mapped to the reference AgamP4 [53, 54] using BWA-MEM v0.7.15 [55] with default settings. Duplicate reads were removed using Sambamba markdup [56]. Freebayes v1.2.0 [57] was used for variant calling, with standard filters and the “-no-population-priors”, “theta = 0.01”, and “max-comple-gap = 0” options. Variants were normalized with *vt normalize* v0.5 [58].

SNPs were filtered out when they did not pass the accessibility mask from Ag1000G, missingness >10%, a minimum depth of 8 and minor allele frequency (MAF) < 1%. In addition, population structure analysis was based on chromosome 3 SNPs only. This was done to avoid confounding signals from polymorphic inversions on chromosomes 2 and X [53]. Heterochromatic regions on chromosome 3R (3R:38,988,757-41,860,198; 3R:52,161,877-53,200,684) and 3L (3L:1-1,815,119; 3L:4,264,713-5,031,692) were also filtered out [53].

Description of population structure was performed by Principal component analysis (PCA) after pruning for LD using scikit-allel v1.2.0 [59]. Hudson’s estimator [60, 61] was used for pairwise fixation indices *F_ST_* calculation implemented in scikit-allel v1.2.0. Nucleotide diversity (π) was calculated in nonoverlapping windows of 10 kb on euchromatic regions of chromosome 3 using VCFtools [62]. The results were grouped by population and significance tests performed between the islands and mainland populations using a Wilcoxon rank-sum test in R.

### (e) Anthropogenic sources of dispersal

The prospects for a GEM emigrating out of a field trial site into a non-target site or vice versa by air or ship transport was assessed by determining the frequency of departures from select mainland and island sites. Airline flight data including the annual (Jan. 1-Dec. 31, 2019) number of international departures from airports within a specific country were obtained from CIRIUM, an aviation data analytics provider [63]. Similarly, shipping data for the annual (Jan.1-Dec.31, 2020) number of commercial ship departures was obtained from the [64].

### (f) Anopheline species richness

Published compilations of Afrotropical *Anopheles* species distributions [65, 66] were used to assemble the information for mainland and island countries. The first criterion for field site selection is the presence of the target species, which is in our case *Anopheles gambiae sensu stricto* and/or its sister species *Anopheles coluzzii*. Species are designated as primary or secondary vectors or as “other” if they are non-vectors or their status as vectors is not clear. Species that commonly had sporozoite infection rates above 1%, as determined by salivary gland dissections, CSP ELISA or PCR of head and thorax were listed as primary vectors. Species with infection rates of <1% were listed as secondary vectors. Our knowledge of the population structure and biology of almost all the secondary vectors is limited and their role in malaria transmission varies from location to location.

## 3. Results and discussion

### (a) Identification of potential field sites

We evaluated 22 potential field sites, including 5 individual islands, multiple islands within 7 archipelagos and 4 islands within Lake Victoria (Figure 1). The sites identified include three island types: continental, oceanic, and lacustrine. Each type possesses features that impact its utility as a GEM trial site. Continental or land bridge islands are unsubmerged portions of the continental shelf and were, at one time, connected to the mainland. Oceanic islands arise from the ocean floor and were never connected to the mainland. Lacustrine islands are islands within lakes and are typically formed by deposits of sedimentary rock, as are the Lake Victoria islands. For comparison, our analyses include mainland sites closest to the islands and those in which GEM field trials are currently under consideration (e.g., Burkina Faso, Mali, Uganda). We then proceed by defining and justifying a prioritized set of criteria (Box 1) on which to base evaluations.

### (b) Geographic isolation

Geographic isolation is among the most significant features favoring islands as GEM field trial sites. Although some mosquito species are known to disperse on prevailing winds over long distances [67, 68], there are, to our knowledge, no reliable reports of open-ocean wind dispersal of malaria vector species over the distances (hundreds of kilometers) separating some of the oceanic islands under consideration here. Emigration of GEMs out of the field trial site into neighboring, non-target sites, either on nearby islands or the mainland, pose a problem, especially as it relates to risk and regulatory concerns. Equally important is immigration of wild type individuals from neighboring sites into the trial site. Immigration, in this case, will confound efforts to measure GEM invasiveness and could potentially render the gene drive inefficient or even ineffective. Island biogeography theory predicts that choosing a remote island as an initial field trial site greatly reduces the potential for gene flow between vector populations both into and out of the island site. This is further supported by the results of our population genomics assessment, as discussed below.

We evaluated geographic isolation for all candidate islands using distance to mainland, UNEP Isolation Index, and SLMP (Table 1). We excluded any island with a UNEP Isolation Index of less than 15 and we used a Surrounding Landmass Proportion (SLMP) value of 1 as a cutoff, so islands with an SLMP value >1 were considered unacceptable. Sites considered unacceptable based on these criteria include the Bijago Islands, Bugala, Koome, Mfangano, Pemba, Zanzibar, and Mafia.

### (c) Island size and topography

There are no well-defined criteria to guide decisions with respect to an appropriately sized area for a GEM field trial. One important consideration is mosquito flight range. To evaluate the dispersal capacity of a GEM, the site should exceed the flight range of the target species. For our considerations we assumed a maximal daily flight range of 10 km for *A. gambiae* [69]. Generally, we aimed to identify sites small enough to be manageable, but large enough to be convincing, keeping the following considerations as a guide.

Area (km^2^) is an important parameter influencing the biology of populations residing on an island. Large island areas typically include more habitat types and can support larger populations. This characteristic can increase the rate of speciation and lower extinction rates over time [27]. Using island size as a criterion we exclude the islands of Annobón and Île Europa for being too small and Madagascar for being too large.

Evaluating the dispersal capabilities of a GEM is a critical outcome from a field trial. This capacity is best evaluated at a site that possess topographical features that may pose a challenge to dispersal, as would be encountered in continental Africa. Elevation was used as a measure of topographic complexity and as a proxy for environmental heterogeneity. The difference between the elevation maximum and minimum of each island measured from sea level is reported in the “Elevation” column in Table 1. Elevation relates to the number of available habitats because of differences between windward and leeward sites, temperature decrease with altitude, and high precipitation regimes at certain altitudes [28].

Altitude and magnitude of steepest gradient were used to generate a graphic representation of topography for each island. A representative sample of these analyses for the islands of Grande Comore and São Tomé are presented in Supplemental Figure 1A and B to illustrate sites having a complex topography and for the islands of Zanzibar and Mafia in Supplemental Figure 1C and D to illustrate a lack of topographic complexity. Sites lacking topographic complexity were excluded from consideration, these included the Bijago Islands, the islands in Lake Victoria, Zanzibar, Pemba, Mafia and Ile Europa.

### (d) Genetic isolation

Genetic isolation relates to the level of gene flow between populations and may be inferred by measuring the degree of genetic divergence between populations under the assumption that gene flow reduces genetic divergence.

Single nucleotide polymorphism (SNP) data were analyzed to reveal genetic relationships among populations and results were visualized using principal components analysis (PCA). The position of individuals in the space defined by the principal components can be interpreted as revealing levels of genetic similarity/dissimilarity among the populations from which those individuals were sampled. Populations occupying the same space are presumed to be very similar genetically and those widely separated, very different.

Results of the PCA for *A. gambiae* populations are illustrated in Figure 3A. This analysis reveals a high degree of genetic similarity between mainland and both lacustrine and continental islands. Conversely, oceanic islands (Comoros archipelago and Madagascar) form discrete individual clusters, indicating that they are genetically distinct both from the mainland and from each other.

**Figure 3.**
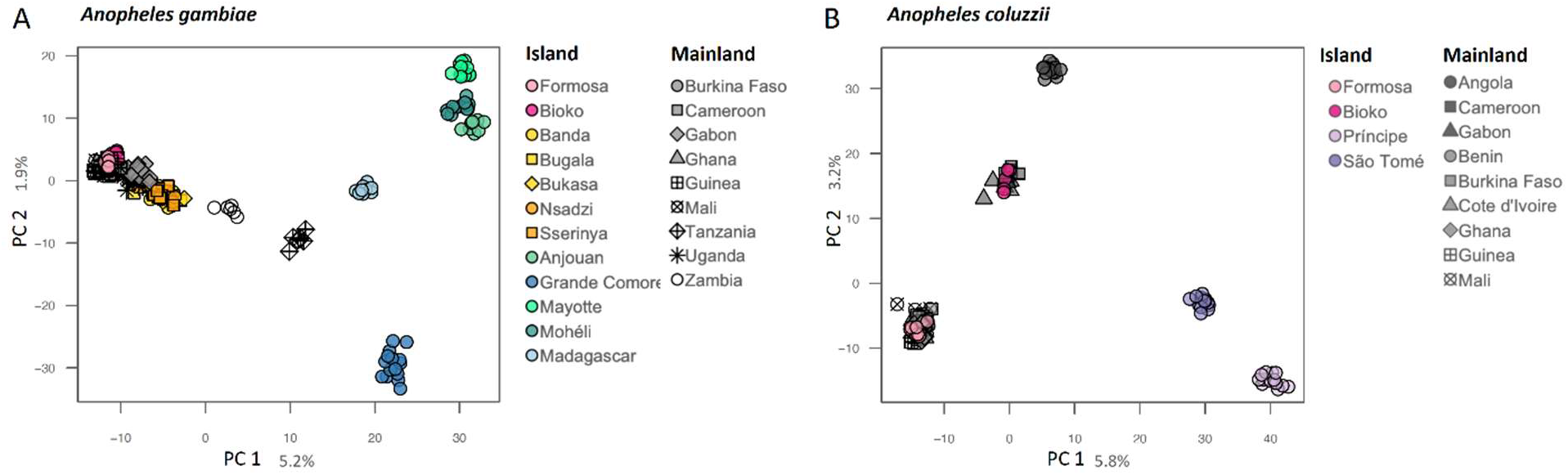
Population structure analysis by PCA. 2D-plot of *A. gambiae* (A) and *A. coluzzii* (B) from islands and mainland populations across Africa. Analyses were based on 50,000 biallelic SNPs from euchromatic regions on chromosome 3. Each marker represents one individual mosquito. Geographic location for each site and numbers of genome analysed per site are provided in Figure 2.

Results of the PCA for *A. coluzzii* (Figure 3B) confirm that populations on continental islands form tight clusters that include mainland populations. Oceanic islands form discrete clusters indicating genetic divergence from mainland populations and from each other. These results indicate high levels of genetic isolation for oceanic island populations of both *A. coluzzii* and *A. gambiae*.

The extent to which individuals move (migrate) between two populations can be approximated by measuring the level of genetic divergence between those populations. Migration (m) can be thought of as including the genotypes of the individuals doing the moving and, in this context, migration results in gene flow. Genetic divergence can be described using the statistic F_ST_, which is the genetic variance in a subpopulation (S) relative to the total variance (T). F_ST_ values range between 0 and 1 and are higher when populations are considerably diverged. The relationship between F_ST_ and *m* is complex, but excluding the effects of drift and selection, the more gene flow between two populations the lower the F_ST_ value. All pairwise F_ST_ values for the populations of *A. gambiae* and *A. coluzzii* analyzed in this study (Figure 2) are presented in Supplemental Figure 2.

Pairwise F_ST_ values for 14 populations of *A. gambiae* spanning its range across sub-Saharan Africa are provided in Supplemental Figure 2A. Results are consistent with the PCA (Figure 3A). West-central populations, including the island of Formosa in the Bijago archipelago, are very similar (F_ST_=0.000-0.009). Divergence between Bioko island and the nearest mainland in Cameroon is higher (F_ST_=0.036). The pattern is quite different in east Africa, wherein mainland populations are far more diverged (F_ST_=0.033-0.090). Divergence between the islands in Lake Victoria and the nearest mainland in Uganda are lower (F_ST_=0.003-0.029). Considerably higher divergence is observed between the Comoros islands and the nearest mainland populations in Tanzania (F_ST_ = 0.130-0.169) and between the Comoros and Madagascar (F_ST_=0.126-0.196).

The F_ST_ values for populations of *A. coluzzii* are likewise consistent with the PCA (Figure 3B). Divergence between the continental island of Formosa and the nearest mainland populations in Guinea-Bissau and between the island of Bioko and nearest sites in Cameroon are low (F_ST_=0.015 and 0.022 respectively). Populations of *A. coluzzii* on the oceanic islands of São Tomé and Príncipe were, by far, the most genetically isolated from mainland populations (F_ST_=0.144 and 0.199 respectively). In addition, the two islands were highly diverged from each other (F_ST_=0.130). The islands within Lake Victoria were excluded from consideration because the *A. gambiae* populations residing on them lacked the high level of divergence that would indicate genetic isolation. No genetic data was available for the Canary Islands, Cape Verde, Zanzibar, Pemba, Mafia, Mauritius, Réunion or Ile Europa.

Taken together, the data summarized in Figures 4 and Supplemental Figure 2 reveal a high degree of genetic isolation among oceanic islands compared with either continental or lacustrine islands. These results suggest limited dispersal (gene flow) between islands and nearest landmasses and are consistent with expectations based on island biogeography theory as described above and reinforce the benefits of selecting a contained island site for conducting GEM field trials. Genetic data is not currently available for several potential island sites, including the Canary Islands, Cape Verde, Île Europa, Zanzibar, Pemba, and Mafia. Genetic isolation, as measured here, was deemed inadequate for the Lake Victoria islands (Bugala, Koome, Mfangano, and Ukara).

**Figure 4.**
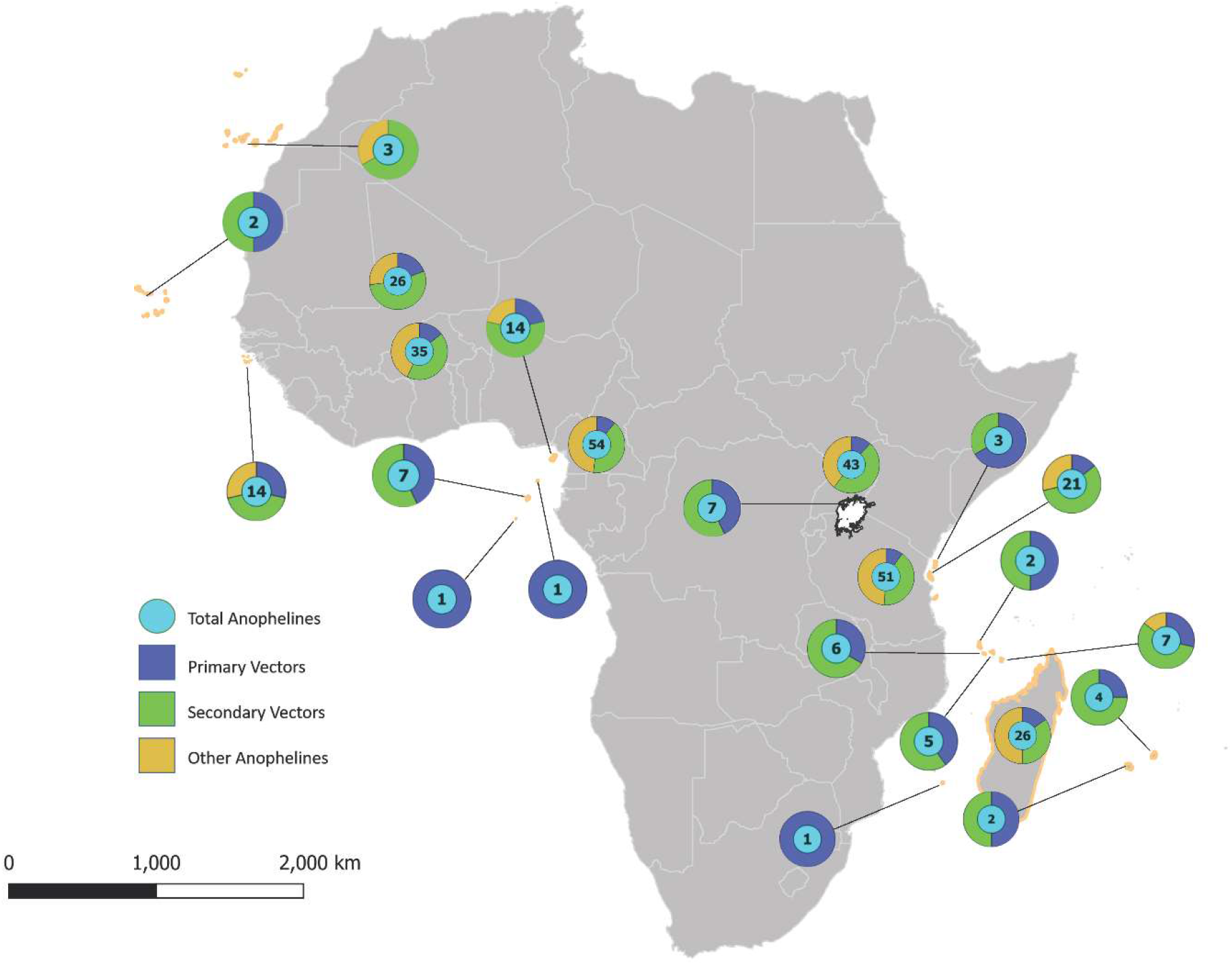
*Anopheles* species complexity in Africa including island and select mainland sites. Map locations and summary of data presented in Table 2. Cyan circle = total number of *Anopheles* spp.; blue proportion of primary vector species; green = proportion of secondary vectors; yellow = proportion of species identified as non-vector or for which vector status unknown.

### (e) Anthropogenic dispersal

Anthropogenic dispersal of mosquitoes from inside the release site into nontarget populations may occur and should be considered in selecting a field trial site. The level of genetic divergence between island and mainland populations of *A. coluzzii* and *A. gambiae* is generally high suggesting that dispersal off the islands is low. Nonetheless, dispersal that may occur is most likely to rely on anthropogenic conveyance [68, 70].

The most significant source of passive anthropogenic dispersal of mosquitoes is by rail and road. This poses significant risk for mainland field sites, where extensive in-country and trans-boundary connections exist [71–73]. Risk by this mode of mosquito dispersal is reduced to zero for oceanic island test sites.

Frequency of air and sea departure to interim and final destinations for a sample of mainland and island populations is presented in Figure 5 and Supplemental Tables 3 and 4. Islands, due to their smaller human populations and geographic areas generally originate less trans-boundary air and sea traffic compared with the continent (Figure 5A). This results in remote islands having inherently lower risk levels for these modes of anthropogenic dispersal. A notable exception is the Cape Verde archipelago which has relatively high ship travel due to its location as a major refueling site (Figure 5B). Traffic has increased with the completion of two new ports and upgrades to existing ports in 1997. Airline and shipping traffic data were only obtained for the locations shown in Figure 5, therefore assessment of the potential for anthropogenic dispersal for the majority of island sites was not assessed. Results for São Tomé and Príncipe and for the Comoros suggest that the likelihood of mosquitoes migrating into or out of these islands is minimal.

**Figure 5.**
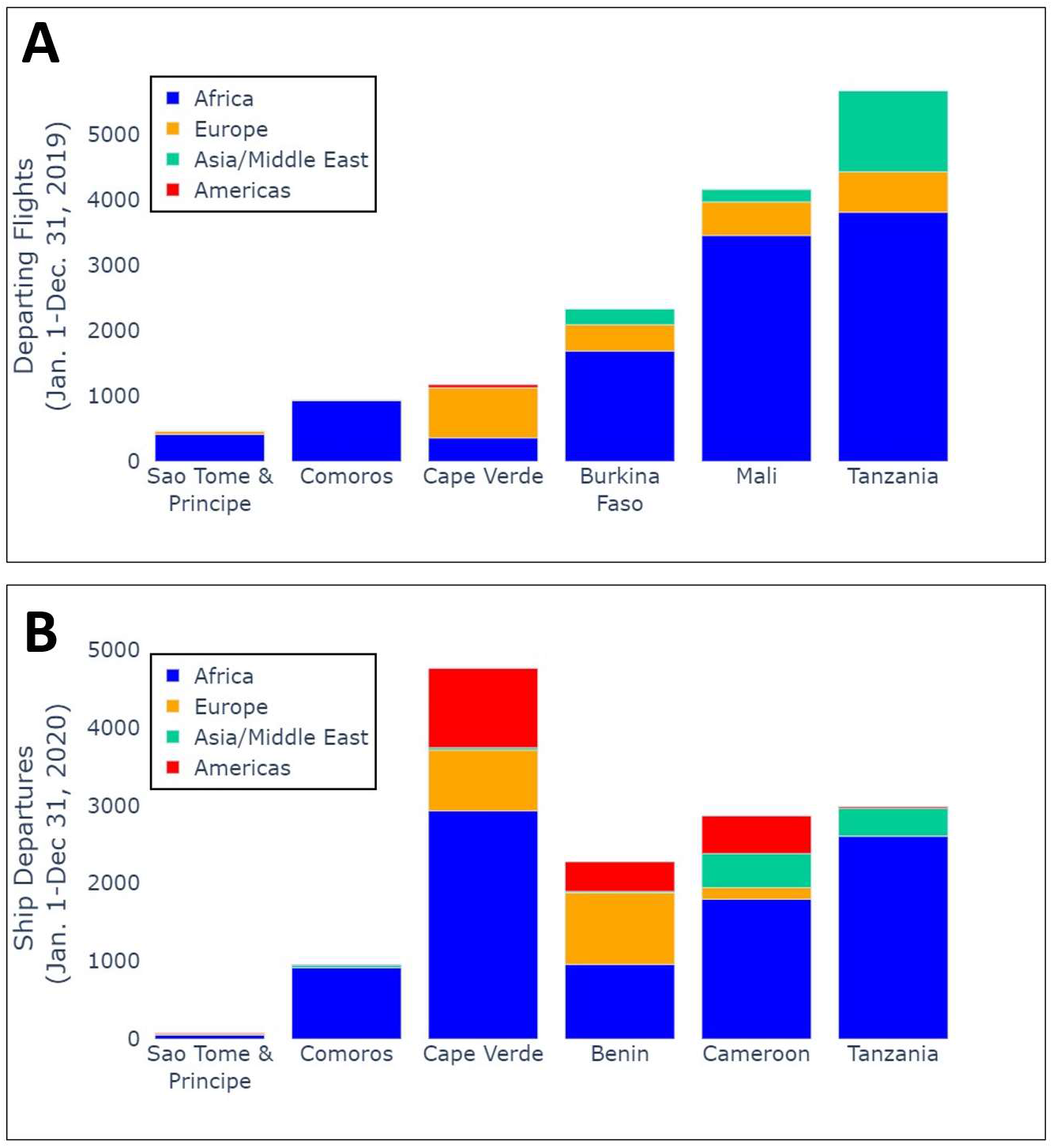
Annual departures by air (A) and sea (B) from representative island and mainland locations in Africa. Colors indicate destinations, grouped by geographic region. Air traffic data provided by Cirium*. (*\*This information has been extracted from a Cirium product. Cirium has not seen or reviewed any conclusions, recommendations or other views that may appear in this document. Cirium makes no warrantees, express or implied, as to the accuracy, adequacy, timeliness, or completeness of its data or its fitness for any particular purpose. Cirium disclaims any and all liability relating to or arising out of use of its data and other content or to the fullest extent permissible by law.*) Sea traffic data provided by the MarineTraffic Global Ship Tracking Intelligence database.

### (f) Anopheline species richness

The number of primary, secondary and other (malaria vector status unclear) species present in island sites and select locations on the mainland are illustrated in Figure 4 (and Supplemental Table 2). It is generally agreed that potential field sites with the fewest number of non-target *Anopheles* species are desirable [16, 74]. If multiple sister species are present, there exists the possibility that the transgene will move between species via natural hybridization [75, 76] which could add an additional level of complexity to post-release assessments. Although the movement of transgene elements between malaria vector species may be considered desirable, it raises the specter of horizontal transfer, which is generally identified as a risk to this technology [77].

Assessment of entomological endpoints following a GEM release requires repeated mosquito collections to quantify changes in the ratio of GEM to wild type mosquitoes. This necessitates sorting large numbers of individual field-collected mosquito samples to separate target from non-target species. For members of sibling species complexes, which are morphologically indistinguishable, this requires the application of PCR-based diagnostics to each individual specimen. If collections include larvae, time-consuming microscopic examination to identify species is required even for those species distinguishable morphologically in the adult stage. Logistically these procedures are greatly simplified where fewer non-target *Anopheles* species are present, positively impacting the time and resources required for successful assessment.

Although entomological endpoints are the main consideration in evaluating the outcome of a PHASE 2 trial, epidemiological impacts should be considered where feasible. If epidemiological endpoints are to be assessed, the presence of multiple primary and secondary vectors are problematic as they can lengthen the season of malaria transmission [78]. Therefore, their presence can mask the effects that GEMs might have on transmission at a field site by maintaining the rate of transmission, even if the parasite is not present in the target mosquito species. If a site is selected in which very few malaria vector species occur, it becomes more likely that the GEM release will have a measurable impact on the level of malaria transmission.

The number of anopheline species present in the mainland sites included here ranged from 26-54. Continental sites (Bijago Islands, Bioko and Zanzibar) had between 14 and 21 species. As expected, oceanic islands contained far fewer, ranging from 1 to 7 species. These results favor the selection of the oceanic islands, São Tomé and Príncipe, Annobón and the Comoros, for field trials. The oceanic islands including the Canary Islands, Cape Verde, Mauritius and Reunion likewise had low numbers of anopheline species, but these were excluded because our target species *A. coluzzii* and *A. gambiae* are absent from these islands.

### (g) Genetic complexity

Genetic complexity was measured using the nucleotide diversity statistic (π), defined as the average number of pairwise nucleotide differences per nucleotide site. Mean nucleotide diversities (π) on oceanic islands (*A. gambiae*- 0.80%, *A. coluzzii*- 0.88%) were significantly lower (p < 0.0001) than mainland population means (*A. gambiae*- 1.17%, *A. coluzzii*- 1.11%) for both species (Figure 6A and B).

**Figure 6.**
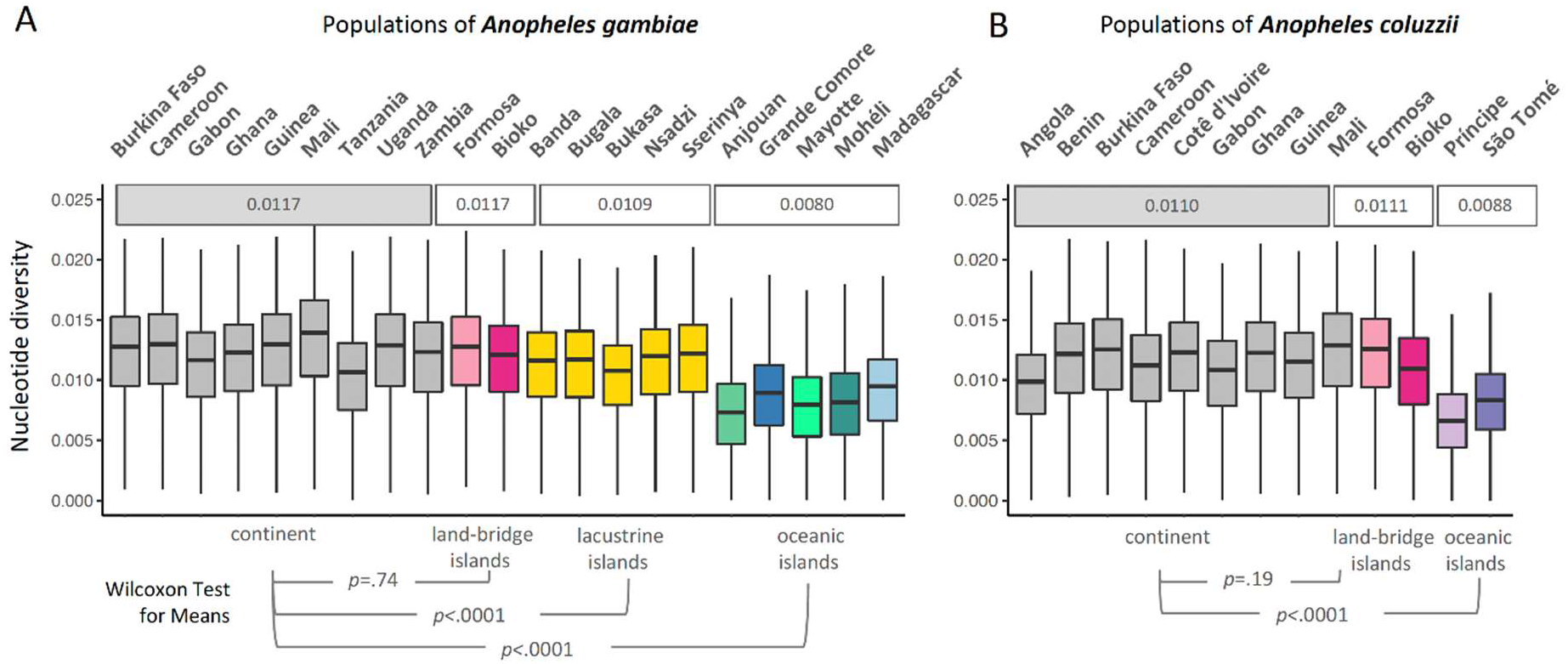
Population diversity. Metric is grouped by sampling locations of (A) *A. gambiae* and (B) *A. coluzzii* populations from island and mainland (grey boxplots). Boxplot of nucleotide diversity (π) performed in 10 kb windows of euchromatic regions of chromosome 3. The midline in all boxplots represents the median, with upper (75th percentile) and lower (25th percentile) limits, whiskers show maximum and minimum values, and outliers are not shown. Mean nucleotide diversity for set of populations are shown above the boxplots; *A. gambiae* populations were divided into four groups: mainland continental (grey), land-bridge (pink), lacustrine (yellow) and oceanic (green/blue) islands; *A. coluzzii* in three: mainland continental (grey), land-bridge (pink) and oceanic (blue) islands. P-value for testing of means between islands and mainland are shown below. Geographic location for each site and numbers of genome analysed per site are provided in Figure 2.

Comparisons among island types yielded results that were consistent with island biogeography theory. Nucleotide diversity in continental island populations did not differ from mainland populations, and lacustrine islands had only slightly lower, but statistically significant, values for π. These observations are expected given the geologic history and proximity of continental and lacustrine islands to the coast. Anjouan island populations presented the lowest (0.73%) nucleotide diversity (π) for *A. gambiae* and Príncipe island for A. coluzzii (0.66%), likely due their small size and high degree of isolation.

In general, the lower biocomplexity on isolated islands includes reduced genetic variation [24]. Our results are concordant with this observation (Figure 6). Selecting field sites with populations containing the lowest levels of variation should decrease the potential for transgene/genome interactions that might negatively impact GEM performance. These include São Tomé and Príncipe and the Comoros.

### (h) Selection of candidate field sites

Each potential site was evaluated based on the criteria listed in Box 1. Evaluations were based on information available from the literature or calculated by us as summarized in the narrative above. Sites that fail to meet all primary criteria were eliminated from further consideration. Those sites that met all primary criteria were raised from potential status to candidate status. Some criteria require further analysis or site visits before a final evaluation can be completed. Sites visits are recommended for candidate sites only. Evaluation of insecticide resistance should be conducted during site visits. Security at candidate sites is dynamic and should be evaluated regularly before and during trials. Evaluation of potential impacts on endangered species requires knowledge about the extent to which these overlap ecologically with A. coluzzii and/or A. gambiae which can only be thoroughly evaluated by mosquito collections made during early site visits.

Overall evaluations are presented in Box 2.

#### BOX 2. Overall summary of evaluation of potential island sites. ✔ =site meets criterion; x=site fails to meet criterion; - = data not available; * = to be determined when site is visited.

| **BOX 2.** Overall summary of evaluation of potential island sites. ✓ =site meets criterion; **x**=site fails to meet criterion; - = data not available; * = to be determined when site is visited. |  |  |  |  |  |  |  |  |  |  |  |
| --- | --- | --- | --- | --- | --- | --- | --- | --- | --- | --- | --- |
| ISLAND | Primary criteria |  |  |  |  |  |  | Other considerations |  |  |  |
|  | A. coluzzii/A. gambiae present | Geographic isolation | Genetic isolation | Genetic diversity | Size (area) | Topography | Anopheline species richness | Insecticide resistance | Plasmodium prevalence | Endangered species | Travel feasibility |
| **OCEANIC ISLANDS** |  |  |  |  |  |  |  |  |  |  |  |
| Canary Islands | **X** | ✓ | - | - | ✓ | ✓ | ✓ | * | **X** | * | ✓ |
| Cape Verde | **X** | ✓ | - | - | ✓ | ✓ | ✓ | * | **X** | * | ✓ |
| Annobón | ✓ | ✓ | ✓ | - | **X** | ✓ | ✓ | * | ✓ | * | **X** |
| São Tomé | ✓ | ✓ | ✓ | ✓ | ✓ | ✓ | ✓ | * | ✓ | * | ✓ |
| Príncipe | ✓ | ✓ | ✓ | ✓ | ✓ | ✓ | ✓ | * | ✓ | * | ✓ |
| Grand Comore | ✓ | ✓ | ✓ | ✓ | ✓ | ✓ | ✓ | * | ✓ | * | ✓ |
| Moheli | ✓ | ✓ | ✓ | ✓ | ✓ | ✓ | ✓ | * | ✓ | * | ✓ |
| Anjouan | ✓ | ✓ | ✓ | ✓ | ✓ | ✓ | ✓ | * | ✓ | * | ✓ |
| Mayotte | ✓ | ✓ | ✓ | ✓ | ✓ | ✓ | ✓ | * | ✓ | * | ✓ |
| Ile Europa | - | ✓ | - | - | **X** | **X** | ✓ | * | - | * | **X** |
| Madagascar | ✓ | ✓ | ✓ | ✓ | **X** | ✓ | **X** | * | ✓ | * | ✓ |
| Mauritius | **X** | ✓ | - | - | ✓ | ✓ | ✓ | * | **X** | * | ✓ |
| Réunion | **X** | ✓ | - | - | ✓ | ✓ | ✓ | * | **X** | * | ✓ |
| **CONTINENTAL ISLANDS** |  |  |  |  |  |  |  |  |  |  |  |
| Bijagos | ✓ | **X** | ✓ | **X** | ✓ | **X** | **X** | * | ✓ | * | ✓ |
| Bioko | ✓ | ✓ | ✓ | - | ✓ | ✓ | **X** | * | ✓ | * | ✓ |
| Zanzibar | ✓ | **X** | - | - | ✓ | **X** | **X** | * | ✓ | * | ✓ |
| Pemba | ✓ | **X** | - | - | ✓ | **X** | ✓ | * | ✓ | * | ✓ |
| Mafia | ✓ | **X** | - | - | ✓ | **X** | - | * | ✓ | * | ✓ |
| **LACUSTRINE ISLANDS (Lake Victoria)** |  |  |  |  |  |  |  |  |  |  |  |
| Bugala | ✓ | **X** | **X** | **X** | ✓ | **X** | ✓ | * | ✓ | * | ✓ |
| Koome | ✓ | **X** | **X** | **X** | ✓ | **X** | ✓ | * | ✓ | * | ✓ |
| Mfangano | ✓ | **X** | **X** | **X** | ✓ | **X** | ✓ | * | ✓ | * | ✓ |
| Ukara | ✓ | **X** | **X** | **X** | ✓ | **X** | ✓ | * | ✓ | * | ✓ |

Evaluation of all twenty-two potential field sites indicate that Bioko, São Tomé & Príncipe, and the Comoros Islands (Anjouan, Grand Comore, Mayotte and Moheli) can be elevated from “potential” to “candidate” GEM field trial sites. The Mascarene (Mauritius and Réunion) and Cape Verde Islands fit many criteria, but Anopheles gambiae does not occur in these islands. Annobón scores well based on several our criteria but travel there was determined to be infeasible, and the island was deemed too small to represent a trial which would provide compelling outcomes.

Therefore, we propose the following as the lead candidate sites for a PHASE 2 GEM field trial: the Comoros Islands, São Tomé and Príncipe and Bioko.

## 4. Conclusions

Our early decision to consider physical islands as the ideal sites for a GEM field trial was guided by contemporary island biogeography theory. This theory provides the basis for certain expectations concerning species richness, in our case, anopheline species richness and also features such as genetic isolation and variability. Our results confirm the relationships between geographic isolation and both genetic isolation (pairwise F_ST_ values) and genetic diversity (nucleotide diversity, pi) which are significantly correlated (Supplemental Figure 3).

The framework described here has been applied by the University of California Irvine Malaria Initiative (UCIMI) as they enter PHASE 2 of GEM research. It is our belief that this comprehensive framework provides identification of site(s) that will maximize the prospect for success, minimize risk, and will serve as a fair, valid, and convincing test of the efficacy and impacts of the UCIMI GEM product, meeting the goal of a PHASE 2 field trial. Furthermore, this process provides a well-reasoned, science-based justification for selecting these sites for GEM field trials, and a solid foundation on which to approach ethical, social, and legal considerations with field site stakeholders.

## Notes

### Competing Interest Statement

The authors have declared no competing interest.

