## Supplementary figures and images for "Selection of Sites for Field Trials of Genetically Engineered Mosquitoes with Gene Drive"

### Supplemental Figure 1

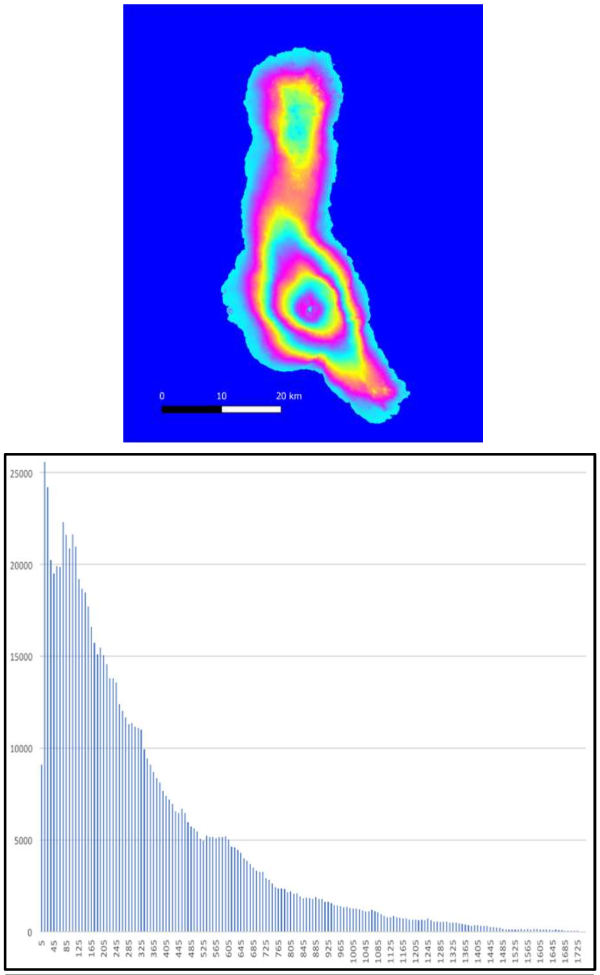


A. Grand Comore topography

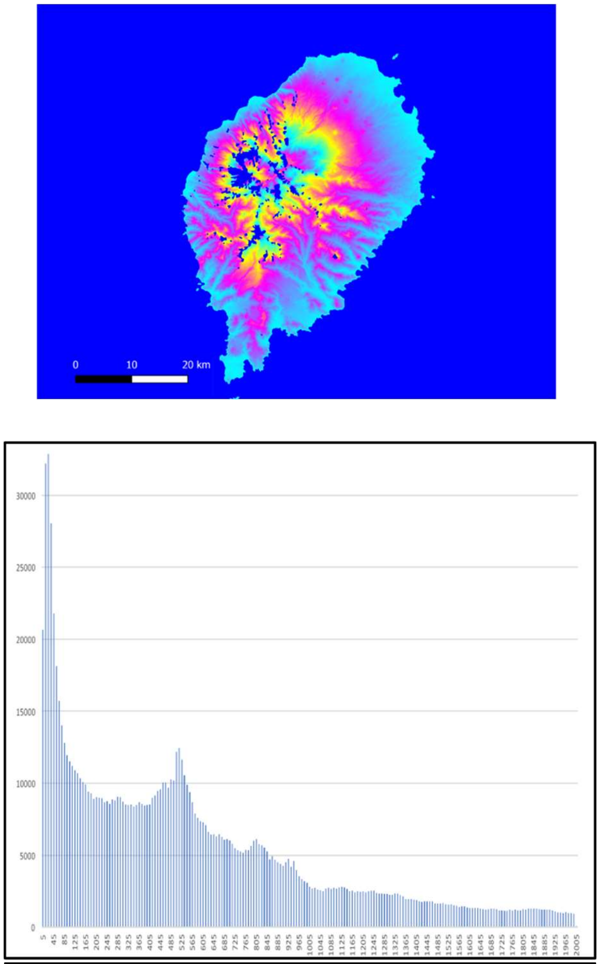


B. São Tomé topography

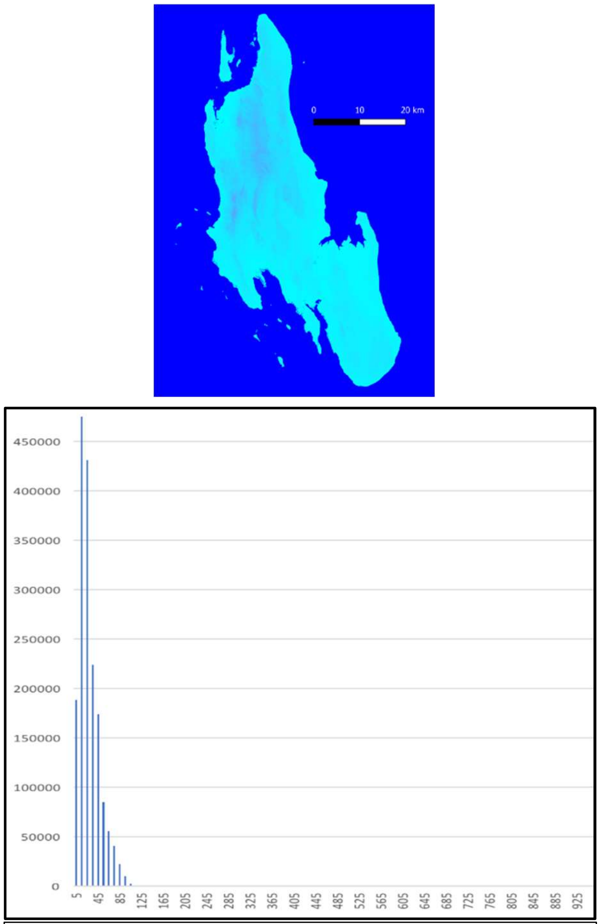


C. Zanzibar topography

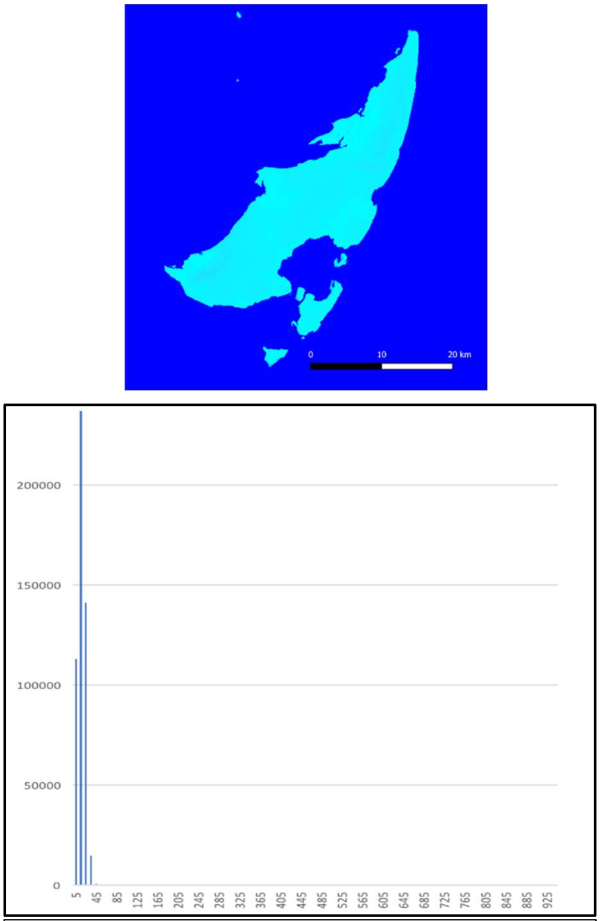


D. Mafia topography

### Supplemental Figure 3

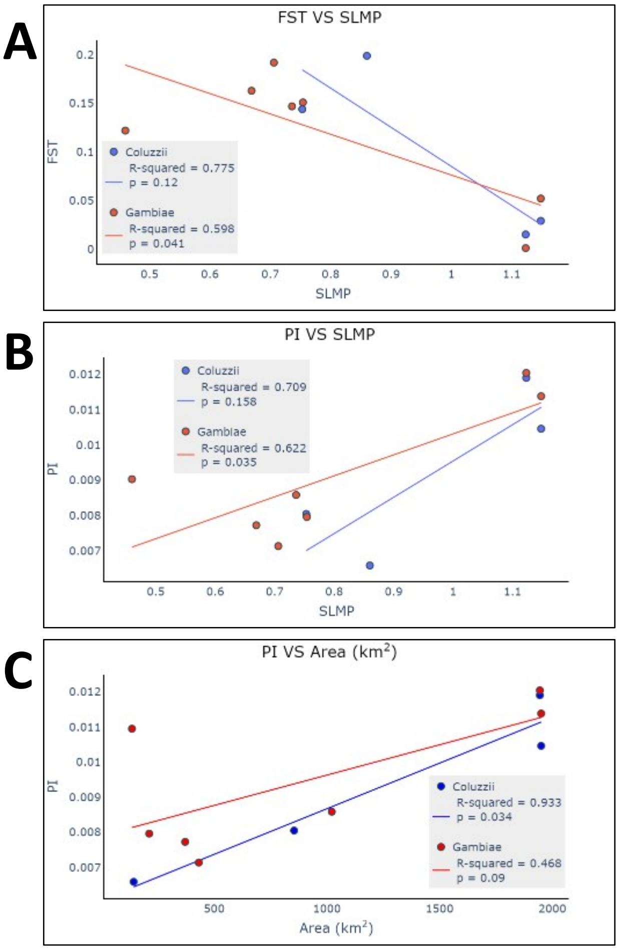
