## Supplemental Figure 2 for "Selection of Sites for Field Trials of Genetically Engineered Mosquitoes with Gene Drive"

A

***Anopheles gambiae*** populations

| | | Zambia | Uganda | The Gambia* | Tanzania | Mali | Guinea-Bissau* | Guinea | Ghana | Gabon | Cameroon | Burkina Faso | Madagascar | Mohéli | Mayotte | Grande Comore | Anjouan | Sserinya | Nsadzi | Bukasa | Bugala | Banda | Bioko |
|---|---|---|---|---|---|---|---|---|---|---|---|---|---|---|---|---|---|---|---|---|---|---|---|
| Island | Formosa | 0.079 | 0.062 | 0.012 | 0.137 | 0.049 | 0.001 | 0.058 | 0.060 | 0.093 | 0.059 | 0.058 | 0.167 | 0.192 | 0.210 | 0.182 | 0.227 | 0.072 | 0.075 | 0.074 | 0.063 | 0.085 | 0.079 |
| Island | Bioko | 0.062 | 0.041 | 0.073 | 0.120 | 0.031 | 0.083 | 0.034 | 0.040 | 0.069 | 0.034 | 0.036 | 0.153 | 0.180 | 0.200 | 0.168 | 0.220 | 0.051 | 0.055 | 0.055 | 0.043 | 0.067 | |
| Island | Banda | 0.042 | 0.029 | 0.078 | 0.099 | 0.037 | 0.089 | 0.042 | 0.046 | 0.075 | 0.041 | 0.043 | 0.133 | 0.162 | 0.180 | 0.146 | 0.203 | 0.020 | 0.029 | 0.028 | 0.022 | | |
| Island | Bugala | 0.024 | 0.003 | 0.055 | 0.081 | 0.014 | 0.066 | 0.016 | 0.022 | 0.052 | 0.016 | 0.018 | 0.118 | 0.146 | 0.164 | 0.130 | 0.186 | 0.010 | 0.014 | 0.012 | | | |
| Island | Bukasa | 0.027 | 0.020 | 0.067 | 0.084 | 0.027 | 0.077 | 0.031 | 0.036 | 0.063 | 0.030 | 0.032 | 0.119 | 0.148 | 0.166 | 0.132 | 0.188 | 0.019 | 0.020 | | | | |
| Island | Nsadzi | 0.033 | 0.019 | 0.067 | 0.088 | 0.027 | 0.077 | 0.031 | 0.037 | 0.064 | 0.031 | 0.032 | 0.122 | 0.153 | 0.171 | 0.134 | 0.192 | 0.019 | | | | | |
| Island | Sserinya | 0.031 | 0.014 | 0.064 | 0.087 | 0.023 | 0.075 | 0.026 | 0.031 | 0.060 | 0.025 | 0.028 | 0.123 | 0.152 | 0.170 | 0.136 | 0.194 | | | | | | |
| Island | Anjouan | 0.164 | 0.195 | 0.224 | 0.191 | 0.182 | 0.230 | 0.204 | 0.208 | 0.223 | 0.204 | 0.204 | 0.193 | 0.137 | 0.174 | 0.201 | | | | | | | |
| Island | Grande Comore | 0.108 | 0.138 | 0.176 | 0.130 | 0.134 | 0.184 | 0.149 | 0.154 | 0.169 | 0.148 | 0.150 | 0.163 | 0.166 | 0.196 | | | | | | | | |
| Island | Mayotte | 0.142 | 0.173 | 0.204 | 0.169 | 0.164 | 0.212 | 0.182 | 0.185 | 0.200 | 0.182 | 0.182 | 0.157 | 0.126 | | | | | | | | | |
| Island | Mohéli | 0.122 | 0.154 | 0.187 | 0.148 | 0.147 | 0.194 | 0.164 | 0.168 | 0.181 | 0.163 | 0.164 | 0.153 | | | | | | | | | | |
| Island | Madagascar | 0.093 | 0.126 | 0.161 | 0.122 | 0.122 | 0.169 | 0.136 | 0.139 | 0.154 | 0.135 | 0.136 | | | | | | | | | | | |
| Mainland | Burkina Faso | 0.043 | 0.016 | 0.050 | 0.101 | 0.002 | 0.063 | 0.000 | 0.008 | 0.052 | 0.003 | | | | | | | | | | | | |
| Mainland | Cameroon | 0.042 | 0.013 | 0.051 | 0.099 | 0.002 | 0.063 | 0.002 | 0.009 | 0.050 | | | | | | | | | | | | | |
| Mainland | Gabon | 0.062 | 0.053 | 0.086 | 0.121 | 0.046 | 0.096 | 0.050 | 0.056 | | | | | | | | | | | | | | |
| Mainland | Ghana | 0.047 | 0.019 | 0.053 | 0.105 | 0.007 | 0.065 | 0.007 | | | | | | | | | | | | | | | |
| Mainland | Guinea | 0.043 | 0.015 | 0.049 | 0.100 | 0.001 | 0.062 | | | | | | | | | | | | | | | | |
| Mainland | Guinea-Bissau* | 0.081 | 0.066 | 0.011 | 0.140 | 0.052 | | | | | | | | | | | | | | | | | |
| Mainland | Mali | 0.036 | 0.012 | 0.041 | 0.090 | | | | | | | | | | | | | | | | | | |
| Mainland | Tanzania | 0.059 | 0.090 | 0.131 | | | | | | | | | | | | | | | | | | | |
| Mainland | The Gambia* | 0.072 | 0.055 | | | | | | | | | | | | | | | | | | | | |
| Mainland | Uganda | 0.033 | | | | | | | | | | | | | | | | | | | | | |

| | Mean $F_{ST}$ between Neighboring | |
|---|---|---|
| | Nearest Mainland | Nearest Island |
| Formosa | 0.001 | - |
| Bioko | 0.052 | - |
| Banda | 0.029 | 0.025 |
| Bugala | 0.003 | 0.014 |
| Bukasa | 0.020 | 0.020 |
| Nsadzi | 0.019 | 0.021 |
| Sserinya | 0.014 | 0.017 |
| Anjouan | 0.192 | 0.171 |
| Grande Comore | 0.147 | 0.188 |
| Mayotte | 0.163 | 0.165 |
| Mohéli | 0.151 | 0.143 |
| Madagascar | 0.122 | 0.167 |

B

***Anopheles coluzzii*** populations

| | | The Gambia* | Mali | Guinea-Bissau* | Guinea | Ghana | Gabon | Cote d'Ivoire | Cameroon | Burkina Faso | Benin | Angola | São Tomé | Príncipe | Bioko |
|---|---|---|---|---|---|---|---|---|---|---|---|---|---|---|---|
| Island | Formosa | 0.030 | 0.028 | 0.015 | 0.053 | 0.032 | 0.088 | 0.031 | 0.076 | 0.032 | 0.037 | 0.133 | 0.176 | 0.225 | 0.090 |
| Island | Bioko | 0.108 | 0.072 | 0.107 | 0.110 | 0.073 | 0.036 | 0.076 | 0.022 | 0.076 | 0.078 | 0.094 | 0.151 | 0.207 | |
| Island | Príncipe | 0.238 | 0.212 | 0.238 | 0.250 | 0.219 | 0.204 | 0.220 | 0.195 | 0.220 | 0.223 | 0.227 | 0.130 | | |
| Island | São Tomé | 0.191 | 0.161 | 0.190 | 0.200 | 0.167 | 0.148 | 0.168 | 0.139 | 0.168 | 0.171 | 0.168 | | | |
| Mainland | Angola | 0.149 | 0.115 | 0.148 | 0.156 | 0.122 | 0.082 | 0.123 | 0.080 | 0.121 | 0.126 | | | | |
| Mainland | Benin | 0.065 | 0.017 | 0.062 | 0.054 | 0.009 | 0.075 | 0.014 | 0.064 | 0.019 | | | | | |
| Mainland | Burkina Faso | 0.061 | 0.001 | 0.060 | 0.056 | 0.018 | 0.072 | 0.015 | 0.062 | | | | | | |
| Mainland | Cameroon | 0.095 | 0.057 | 0.094 | 0.096 | 0.060 | 0.024 | 0.061 | | | | | | | |
| Mainland | Cote d'Ivoire | 0.059 | 0.014 | 0.054 | 0.045 | 0.008 | 0.074 | | | | | | | | |
| Mainland | Gabon | 0.108 | 0.068 | 0.106 | 0.108 | 0.071 | | | | | | | | | |
| Mainland | Ghana | 0.061 | 0.016 | 0.057 | 0.048 | | | | | | | | | | |
| Mainland | Guinea | 0.079 | 0.051 | 0.071 | | | | | | | | | | | |
| Mainland | Guinea-Bissau* | 0.011 | 0.053 | | | | | | | | | | | | |
| Mainland | Mali | 0.055 | | | | | | | | | | | | | |

| | Mean $F_{ST}$ between Neighboring | |
|---|---|---|
| | Nearest Mainland | Nearest Island |
| Formosa | 0.015 | - |
| Bioko | 0.029 | 0.207 |
| Príncipe | 0.199 | 0.130 |
| São Tomé | 0.144 | 0.130 |
