## Supplemental Figure Legends for "Selection of Sites for Field Trials of Genetically Engineered Mosquitoes with Gene Drive"

**Supplemental Figure 1.** Island topography. Satellite images and elevation histograms of Grande Comore (**A**), São Tomé, (**B**), Zanzibar (**C**) and Mafia (**D**). The satellite images show the relief, or variations in topographical complexity, with warmer colors signifying high elevation points and cooler colors signifying low elevation. Histograms show "Magnitude of steepest gradient" display amount of vertical change per unit of horizontal distance in the direction of maximum change.

**Supplemental Figure 2**. Population structure analysis by $F_{ST}$ analyses. Genetic differentiation between island and mainland African populations of *A. gambiae* **(A)** and *A. coluzzii* **(B)**. Analyses were based on biallelic SNPs on euchromatic regions on chromosome 3. Gray shades highlight low (light) to high (dark) values. Insert tables highlights the mean $F_{ST}$ between each island population and it's geographically proximal mainland and island populations. Geographic location for each site and numbers of genome analysed per site are provided in Figure 4. *population with *A. coluzzii*/*A. gambiae* hybrid individuals.

**Supplemental Figure 3.** A. Shows a negative correlation between the proportion of surrounding land mass (SLMP) of candidate islands plotted against FST of the closest mainland country for island populations where both genetic data and SLMP data are available. B. Shows a positive correlation between the proportion of surrounding land mass (SLMP) of candidate islands plotted against average nucleotide diversity (PI) for island populations where both genetic data and SLMP data are available. C. Shows a positive correlation between the area (km2) of candidate islands plotted against the average nucleotide diversity (PI) for island populations where both genetic data and area data are available. All plots and correlation statistics were generated using the python. Correlation trendlines for An. gambiae and An. coluzzii were calculated separately.
