## Supplemental Table 2 for "Selection of Sites for Field Trials of Genetically Engineered Mosquitoes with Gene Drive"

**Supplemental Table 4.** Ship Traffic

| PORT | COUNTRY | DESTINATION | | | | | | | |
|---|---|---|---|---|---|---|---|---|---|
| | | Africa | Asia | Australia | Europe | Middle East | N. America | Russia | S. America |
| COTONOU | Benin | 959 | 13 | 0 | 921 | 5 | 29 | 4 | 351 |
| DAR ES SALAAM | Tanzania | 2606 | 264 | 2 | 4 | 94 | 5 | 0 | 14 |
| DOUALA | Cameroon | 1796 | 438 | 0 | 150 | 1 | 122 | 0 | 361 |
| DURBAN | South Africa | 2553 | 580 | 54 | 135 | 210 | 219 | 1 | 297 |
| MINDELO | Cape Verde | 1771 | 23 | 2 | 750 | 6 | 613 | 6 | 362 |
| PRAIA | Cape Verde | 1162 | 0 | 4 | 33 | 0 | 13 | 0 | 32 |
| FOMBONI | Comoros | 2 | 0 | 0 | 0 | 0 | 0 | 0 | 0 |
| MORONI | Comoros | 326 | 1 | 0 | 0 | 14 | 9 | 0 | 0 |
| MUTSAMUDU | Comoros | 588 | 0 | 0 | 0 | 18 | 0 | 0 | 0 |
| PRINCIPE | Sao Tome and Principe | 23 | 0 | 0 | 0 | 0 | 0 | 0 | 0 |
| SAO TOME | Sao Tome and Principe | 28 | 1 | 0 | 21 | 0 | 3 | 0 | 3 |
