## Supplemental Table 3 for "Selection of Sites for Field Trials of Genetically Engineered Mosquitoes with Gene Drive"

**Supplemental Table Legends**

**Supplemental Table 1.** Metadata per sample used for population genomics analyses. M = *Anopheles coluzzii*, S=*Anopheles gambiae* s.s., VGL= University of California Davis Vector Genetics Laboratory.

**Supplemental Table 2**. Comparison of mainland versus island Anopheles species diversity and malaria incidence. Species of *Anopheles* present in specific Afrotropical countries and Islands listed according to their malaria vector status (primary vector, secondary vector or other (non-vector/status unclear) also included are malaria case numbers for countries and islands for which information is available. Species names in bold are likely present, but not confirmed.

**Supplemental Table 3.** Airline departures from airports in a sample of mainland and island airports. Data for a one-year period, January 1 through December 31, 2019. ******This information has been extracted from a Cirium product. Cirium has not seen or reviewed any conclusions, recommendations or other views that may appear in this document. Cirium makes no warrantees, express or implied, as to the accuracy, adequacy, timeliness, or completeness of its data or its fitness for any particular purpose. Cirium disclaims any and all liability relating to or arising out of use of its data and other content or to the fullest extent permissible by law.*

**Supplemental Table 4.** Outgoing ship traffic and destinations. Ports across Africa showing the number of ships that left that specific port and the geographic area where those ships traveled to after leaving that port. The data shows that the number of ships departing Sao Tome and Principe and the Comoros is greatly reduced compared to other ports in Africa. Data was obtained from the marinetraffic.com database and covers the one-year period between January 1 to December 31, 2020.
