## Supplemental Table Legends for "Selection of Sites for Field Trials of Genetically Engineered Mosquitoes with Gene Drive"

**Supplemental Table 2.** Comparison of mainland versus island *Anopheles* species diversity and malaria incidence. Species of *Anopheles* present in specific Afrotropical countries and Islands listed according to their malaria vector status (primary vector, secondary vector or other (non-vector/status unclear) also included are malaria case numbers for countries and islands for which information is available. Species names in bold are likely present, but not confirmed.

**MAINLAND**

| Country | Primary Vectors | Secondary Vectors | Other *Anopheles* (non-vector or status unclear) | References | Malaria Cases (per year) |
|---|---|---|---|---|---|
| Burkina Faso | *An. arabiensis, coluzzii, funestus, gambiae, nili* | *An. brunnipes, coustani, cydippis, hancocki, leesoni, maculipalpis, paludis, pharoensis, rivulorum, rufipes, pretoriensis, sergentii, squamosus, theileri, ziemanni* | *An. argenteolobatus, brohieri, brumpti, domicolus, dureni, flavicosta, freetownensis, implexus, longipalpis, murphyi, natalensis, obscurus, rhodesiensis, somalicus, wellcomei* | | 6,840,864 (2000)<br>8,602,187 (2010)<br>7,245,827 (2015)<br>7,859,000 (2019) |
| Cameroon | *An. arabiensis, coluzzii, funestus, gambiae, moucheti, nili* | An. *bervoetsi, brunnipes, carnevalei, coustani, cydippis, demeilloni, hancocki, leesoni, maculipalpis, marshallii, melas, obscurus, ovengensis, paludis, pharoensis, pretoriensis, rivulorum, rivulorum*-like, *rufipes, sergentii, squamosus, ziemanni* | *An. brohieri, buxtoni, christyi, cinctus, concolor, deemingi, domicolus, dualaensis, eouzani, flavicosta, freetownensis, hargreavesi, implexus, jebudensis, longipalpis, mousinhoi, multicinctus, namibiensis, natalensis, okuensis, rageaui, rhodesiensis, smithii, somalicus, tenebrosus, wellcomei* | | 6,291,500 (2000)<br>5,909,335 (2010)<br>5,777,768 (2015)<br>6,291,256 (2019) |
| Mali | *An. arabiensis, coluzzii, funestus, gambiae, nili* | *An. brunnipes, coustani, dthali, hancocki, leesoni, maculipalpis, paludis, pharoensis, pretoriensis, rivulorum, rufipes, sergentii, squamosus, ziemanni* | *An. brohieri, domicolus, flavicosta, obscurus, rhodesiensis, somalicus, wellcomei* | 1,2 | 4,446,769 (2000)<br>5,772,983 (2010)<br>6,833,022 (2015)<br>6,560,000 (2019) |
| Tanzania | *An. arabiensis, funestus, gambiae, moucheti, nili* | *An. aruni, brunnipes, cinereus, coustani, cydippis, demeilloni, gibbinsi, leesoni, maculipalpis, marshallii, merus, paludis, parensis, pharoensis, pretoriensis, quadriannulatus, rivulorum, rufipes, squamosus, theileri, ziemanni* | *An. ardensis, argenteolobatus, christyi, confusus, distinctus, erepens, garnhami, implexus, keniensis, kingi, letabensis, longipalpis, lovettae, machardyi, namibiensis, natalensis, njombiensis, rhodesiensis, schwetzi, seydeli, swahilicus, tenebrosus, walravensi, wellcomei, wilsoni* | | 11,514,222 (2000)<br>5,917,848 (2010)<br>7,298,719 (2015)<br>6,453,096 (2019) |
| Uganda | *An. arabiensis, funestus, gambiae, moucheti, nili* | *An. bwambae, cinereus, coustani, cydippis, demeilloni, gibbinsi, hancocki, leesoni, maculipalpis, marshallii, paludis, parensis, pharoensis, pretoriensis, quadriannulatus, rivulorum, rufipes, squamosus, symesi, theileri, ziemanni* | *An. ardensis, brohieri, christyi, domicolus, garnhami, hargreavesi, harperi, implexus, keniensis, kingi, longipalpis, natalensis, obscurus, rhodesiensis, tenebrosus, vinckei, wellcomei* | | 11,522,961 (2000)<br>13,277,279 (2010)<br>9,690,714 (2015)<br>11,629,246 (2019) |

**ISLANDS**

| Island | Primary Vectors | Secondary Vectors | Other *Anopheles* (non-vector or status unclear) | | Malaria Cases (year) |
|---|---|---|---|---|---|
| Bijagos Islands (Guinea-Bissau) | *An. arabiensis, coluzzii, gambiae* | *An. melas,* ***cinereus, coustani, maculipalpis, pharoensis, rufipes, squamosus, ziemanni*** | ***An. dancailicus, hargreavesi, smithii*** | 3, 4 | |
| Bioko (Equatorial Guinea) | *An. coluzzii, funestus, gambiae,* ***moucheti*** | ***An. brunnipes, carnevalei, leesoni, melas, ovengensis*** | ***An. cinctus, lloreti, obscurus, smithii*** | 5, 6 | |
| Canary Islands (Spain) | | *An.multicolor, sergentii* | *An.hispaniola* (historically a vector in Europe) | 7 | |
| Cape Verde | *An. arabiensis* | *An. pretoriensis* | | 8 | 144 (2000)<br>47 (2010)<br>7 (2015)<br>0 (2019) |
| Anjouan (Comoros) | *An. funestus, gambiae* | *An. coustani, mascarensis, pretoriensis* | | 9, 10, 11 | 35,309 (2000)<br>36,538 (2010)<br>1,300 (2015)<br>17,599 (2019) |
| Grand Comore | *An. gambiae* | *An. pretoriensis* | | | |

| | | | | | |
|---|---|---|---|---|---|
| (Comoros) | | | | | |
| Moheli (Comoros) | *An. funestus, gambiae* | *An. coustani, maculipalpis, mascarensis, pretoriensis* | | | |
| Lake Victoria islands | *An. arabiensis, funestus, gambiae* | *An. coustani, pharoensis, symesi, ziemanni* | | 12, 13, 14 | |
| Mayotte (France) | *An. funestus, gambiae* | *An. coustani, maculipalpis, mascarensis, pretoriensis* | *An. comorensis* | 1 | >2,000 (2000)<br>433 (2010)<br>14 (2015)<br>22 (2016) |
| Île Europa (France) | *An. gambiae* | | | 15 | |
| Madagascar | *An. arabiensis, funestus, gambiae, coustani* | *An. brunnipes, cydippis, maculipalpis, mascarensis, merus, pharoensis, pretoriensis, rufipes, squamosus* | *An. flavicosta, fuscicolor, grassei, grenieri, griveaudi, lacani, milloti, notleyi, pauliani, radama, ranci, roubaudi, tenebrosus* | 1 | 901,335 (2000)<br>893,540 (2010)<br>1,897,533 (2015)<br>2,052,071 (2019) |
| Pemba (Tanzania) | *An. arabiensis, gambiae* | *An. merus* | | 17 | |
| Príncipe | *An. coluzzii* | | | 18, 19 | 31,975 (2000)<br>2,740 (2010)<br>2,058 (2015)<br>2,446 (2019) |
| São Tomé | *An. coluzzii, funestus, gambiae* | *An. coustani, melas, paludis, pharoensis* | | | |
| Annobón | *An. coluzzii* | | | 20 | |
| Mauritius | *An. arabiensis* | *An. coustani, maculipalpis, merus* | | 21 | All imported:<br>52 (2010)<br>33 (2012) |
| Réunion (France) | *An. arabiensis* | *An. coustani* | | 22 | Eliminated (1979) |
| Zanzibar (Tanzania) | *An. arabiensis, funestus, gambiae* | *An. aruni, coustani, leesoni, maculipalpis, marshallii, merus, paludis, parensis, pretoriensis, rivulorum, squamosus, ziemanni* | *An. longipalpis, obscurus, quadriannulatus, swahilicus, tenebrosus, wellcomei* | 2 | 3,528 (2005)<br>2,572 (2012)<br>3,814 (2015)<br>3,025 (2016) |

TABLE 2 REFS:

[10]Brunhes, J., Le Goff, G., & Geoffroy, B. (1997). Anophèles afro-tropicaux : 1. Descriptions d'espèces nouvelles et changements de statuts taxonomiques (Diptera : Culicidae). *Annales de la Societe Entomologique de France, 33*.
